# Clonal determinants of organotropism and survival in metastatic uveal melanoma

**DOI:** 10.1101/2024.05.14.593919

**Authors:** Bailey S.C.L. Jones, Patrick C. Demkowicz, Mitchelle Matesva, Renelle Pointdujour Lim, John H. Sinard, Antonietta Bacchiocchi, Ruth Halaban, Marcus Bosenberg, Mario Sznol, Harriet M. Kluger, Mathieu F. Bakhoum

**Affiliations:** Department of Ophthalmology & Visual Science, Yale School of Medicine, New Haven, Connecticut; Yale School of Medicine, New Haven, Connecticut; Institute of Dermatology & Oculoplastic Surgery, Sarasota, Florida; Department of Pathology, Yale School of Medicine, New Haven, Connecticut; Department of Dermatology, Yale School of Medicine, New Haven, Connecticut; Yale Cancer Center, Yale School of Medicine, New Haven, Connecticut; Department of Medicine, Yale School of Medicine, New Haven, Connecticut

## Abstract

Uveal melanoma (UM), the most common intraocular primary cancer in adults, demonstrates a unique proclivity for liver metastasis. To understand the molecular underpinnings of this organotropism, we analyzed the genomic features of liver and extrahepatic UM metastases, identifying distinct molecular signatures that mirror the clonal diversity in primary UM tumors. Liver metastases were enriched in *BAP1* mutations and exhibited a higher prevalence of monosomy 3 compared to extrahepatic metastases. Analysis of the tumor-liver microenvironment crosstalk at the single-cell level underscored a significant role for hepatic stellate cells in facilitating UM growth and establishment in the liver. Notably, within the primary tumor, clones that demonstrated a high affinity for the liver, compared to those with low liver affinity, exhibited a distinct transcriptional profile characterized by the upregulation of pathways that activate hepatic stellate cells, specifically involving TGF-β signaling, cytokine signaling, extracellular matrix remodeling, and angiogenesis. Liver-tropic clones displayed not only an increased affinity for liver colonization but were also associated with worse survival outcomes, underscoring the adverse prognostic significance of hepatic metastases in UM. Our findings demonstrate that trajectories of metastatic dissemination and patient survival in UM are established early in the primary tumor’s evolution, opening pathways for the development of targeted therapeutic interventions to improve patient outcomes.

## Introduction

Metastasis, the process by which cancer cells spread to distant organs, is influenced by the interplay between cancer cells and their target organ microenvironments. This phenomenon, known as organotropism, has been observed in various malignancies, including uveal melanoma (UM), the most common intraocular cancer in adults^1–3^.The majority (90%) of patients that develop UM metastases have liver involvement, establishing UM as an excellent model for studying organotropism in cancer^4,5^. Notably, patients with UM metastases in the liver experience significantly worse survival outcomes than those with only extrahepatic metastases^6^, emphasizing the need to understand the molecular factors driving liver tropism in UM.

To elucidate tumor-intrinsic factors underlying organotropism in UM, we analyzed whole exome and RNA sequencing of both hepatic and extrahepatic UM metastases, single-cell and single-nucleus RNA sequencing of primary and metastatic specimens, as well as clinical outcomes of patients with primary and metastatic UM. Our findings from single-cell RNA sequencing of primary UM tumors identified two predominant clonal types, each characterized by unique mutational, transcriptional, and epigenetic profiles^7^. Interestingly, while both clonal types demonstrated the capacity to metastasize to extrahepatic organs, hepatic metastases were dominated by one specific clonal type. Analysis of ligand-receptor interactions within the liver metastatic environment underscored a significant role for hepatic stellate cells. Clones from the primary tumor with increased predisposition for liver metastasis exhibited transcriptional programs known to activate stellate cells. These findings suggest that the pattern of metastatic dissemination in UM is not an arbitrary event; instead, it is pre-defined by specific tumor clones that exist in the primary tumor. These insights not only enhance our understanding of organ-specific metastasis patterns in UM but also pave the way for promising opportunities in developing targeted therapies, ultimately improving patient outcomes.

## Results

We first examined the distribution of hallmark UM mutations in metastasis. In primary UMs, there are two clusters of genes with recurring mutations^8^. The first cluster includes the mutually exclusive *GNAQ* and *GNA11*, which are implicated in UM tumorigenesis. The second cluster includes mutations in *BAP1*, *EIF1AX*, and *SF3B1*, which provide prognostic information and exhibit a pattern of mutual exclusivity, although not as pronounced as in the first cluster^9^. In The Cancer Genome Atlas (TCGA)^10^, there were 34 primary UM samples with only *GNAQ* mutations, 38 with only *GNA11* mutations, and 2 with both *GNAQ* and *GNA11*, with a Jaccard index of 0.03, indicating high degree of mutual exclusivity. In the second cluster of genes, there were 12 samples with only *BAP1* mutations, 9 with only *SF3B1*, 9 with only *EIF1AX*, 1 with both *BAP1* and *SF3B1* (Jaccard index 0.04), and 1 with both *BAP1* and *EIF1AX* mutations (Jaccard index 0.05) (Fig. 1a,b). These mutation patterns were also observed in metastatic samples. We analyzed whole exome or whole genome sequencing from 144 metastatic UMs procured at our institution and from published cohorts (refer to Extended Data Table 1 for additional details)^11–14^. There were 61 samples with only *GNAQ* mutations, 69 with only *GNA11*, maintaining a pattern of high mutual exclusivity as in primary tumors. In the second cluster of genes, there were 77 samples with only *BAP1* mutations, 28 with only *SF3B1*, 7 with only *EIF1AX*, 6 with both *BAP1* and *SF3B1* (Jaccard index 0.05), and 3 with both *BAP1* and *EIF1AX* mutations (Jaccard index 0.03) (Fig. 1a,b). The similarity in mutation distribution between primary and metastatic UM suggests a preserved clonal structure between the primary and the metastatic setting.

**Figure 1.**
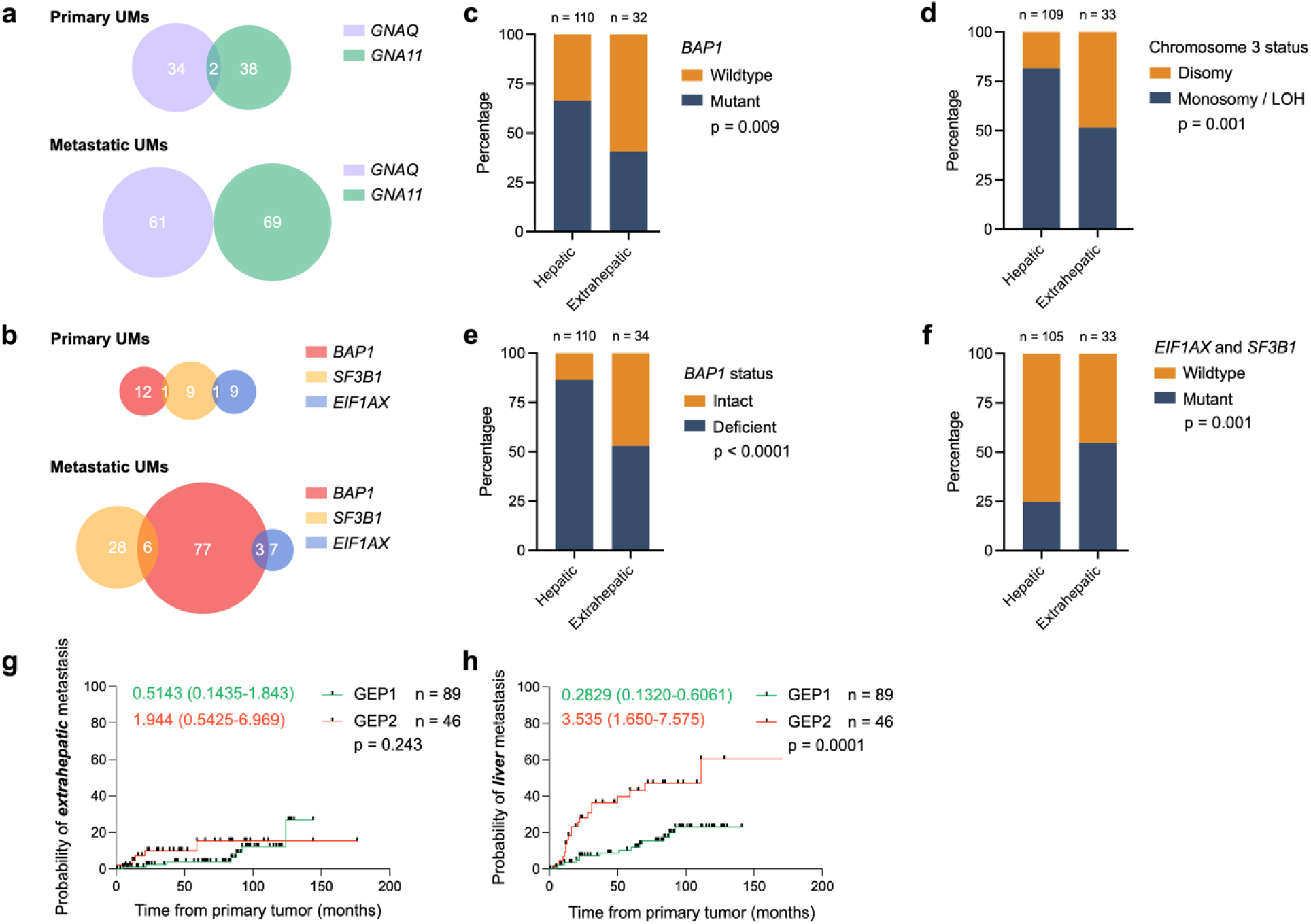
**a**, Proportional Venn diagram of primary UMs with *GNAQ* (purple), *GNA11* (green), *BAP1* (red), *SF3B1* (yellow), and *EIF1AX* (blue) mutations. **b**, Proportional Venn diagram of primary UMs with *GNAQ* (purple), *GNA11* (green), *BAP1* (red), *SF3B1* (yellow), and *EIF1AX* (blue) mutations. Proportion of **c**, *BAP1* mutations (mutant shown in blue and wildtype shown in orange), **d**, chromosome 3 copy number (monosomy 3 or loss of heterozygosity (LOH) shown in blue and disomy 3 shown in orange), **e**, *BAP1* status (deficient (mutant or genomic copy loss) shown in blue and intact shown in orange), **f**, *EIF1AX* and/or *SF3B1* mutations (mutant shown in blue and wildtype shown in orange) in hepatic and extrahepatic metastases. Statistical significance was determined using a two-sided Chi-square test. Kaplan-Meier curve showing the probability of (**g**) hepatic and (**h**) extrahepatic metastasis in patients with primary UM stratified by their bulk gene expression profile (GEP1 shown in green and GEP2 shown in red). Statistical significance was determined using the Log-rank test. *Source: **a***, *Robertson et al.*^10^*; **b**,**c**,**d**,**e**,**f**, Yale, Ny et al.*^13^*, Karlsson et al.*^12^*, Nguyen et al.*^14^*, and Royer-Bertrand et al.*^11^*; **g***,***h****, Yale*

We then examined the distribution of these mutations in hepatic versus extrahepatic metastases and found that liver metastases were enriched in *BAP1* mutations compared to extrahepatic metastases: 66.4% vs. 40.6%, respectively (*P* = 0.009) (Fig. 1c). Given the insights from TCGA data, which suggested that whole exome sequencing does not capture all *BAP1* mutations in primary UMs, especially intronic ones^10^, we also assessed the prevalence of monosomy 3 in liver metastases. Of note, *BAP1* is located on the short arm of chromosome 3, and primary UMs with *BAP1* mutations almost invariably exhibit monosomy 3^9^. We found that 81.7% of liver metastases displayed monosomy 3, in contrast to 51.5% of extrahepatic metastases (*P* = 0.001) (Fig. 1d). Altogether, 86.4% of liver metastases were deficient in *BAP1* due to mutations or genomic copy loss, compared to 52.9% of extrahepatic metastases (*P* < 0.001) (Fig. 1e). Conversely, extrahepatic metastases showed an enrichment for *SF3B1* or *EIF1AX* mutations compared to liver metastases: 54.5% vs. 24.8%, respectively (*P* = 0.001) (Fig. 1f). Furthermore, we found that mutations in *GNAQ* or *GNA11* were equally prevalent in both hepatic and extrahepatic metastases (Extended Data Fig. 1a,b). These findings suggest a selective pressure in hepatic metastases towards a specific clone characterized by monosomy 3, frequent *BAP1* mutations, and wildtype *EIF1AX* or *SF3B1*. This pattern implied that metastatic organotropism in UM is pre-determined by the primary tumor clones’ subtypes, as identified through their hallmark mutations.

To further substantiate the connection between metastatic tropism and the primary tumor clones, we asked whether the phenotypic classification of primary UMs could predict subsequent patterns of metastatic spread. In clinical practice, primary UMs can be stratified into high-risk and low-risk groups based on DNA analysis or gene expression profiling (GEP). Tumors with a more favorable prognosis typically are wildtype for *BAP1*, may harbor *EIF1AX* or *SF3B1* mutations, are disomic for chromosome 3, and have a GEP class 1 signature. Conversely, those with worse prognosis often have *BAP1* mutations, are monosomic for chromosome 3, and exhibit GEP class 2 signature^10,15–17^. With the new insight gained from molecular data linking metastatic tropism to clonal diversity, we examined the incidence of organ-specific metastases in 135 individuals from our institution stratified by GEP classification of their primary tumor samples and with long-term follow-up data (refer to Extended Data Table 2 for additional details). In line with our previous findings that both clonal types have the potential to metastasize to extrahepatic organs, there was no significant difference in the rates of extrahepatic metastasis between GEP class 1 and class 2 tumors (*P* = 0.243, HR = 1.94, 95% CI = 0.543-7.82) (Fig. 1g), whereas individuals with GEP class 2 primary UMs showed a significantly higher propensity for developing liver-specific metastases compared to those with GEP class 1 tumors (*P* = 0.0001, HR = 3.54, 95% CI = 1.65-7.58) (Fig. 1h). This marked disparity in metastatic patterns underscores our hypothesis that the trajectories of metastatic dissemination in UM are pre-determined in the primary tumor.

To identify the cellular interactions that promote the proliferation of UM cells in the liver, we examined ligand-receptor interactions between tumor cells and the liver microenvironment, using data from single-cell RNA sequencing of three metastatic UM hepatic samples^18^ and from single-nucleus RNA sequencing of seven treatment-naïve metastatic UM hepatic samples^19^. Cell identities were validated using cell-specific canonical marker genes. Single-cell RNA sequencing identified various cell populations: melanoma cells (65.9%), immune cells (32.6%), and endothelial cells (1.5%) (Extended Data Fig. 2a,b,c). Single-nucleus RNA sequencing revealed the presence of melanoma cells (72.9%), immune cells (3.6%), endothelial cells (1.0%), hepatocytes (19.5%), and notably, hepatic stellate cells (3.0%) (Extended Data Fig. 2d,e,f). Detection of stellate cells in the single-nucleus RNA sequencing dataset underscores its advantage over single-cell RNA sequencing, particularly in isolating specific liver cell types that are otherwise challenging to capture^20^. We used CellChat^21^, a computational tool designed to infer, analyze, and visualize cell-cell communication from single-cell data to identify ligand-receptor pairs among different cell types. Our analysis revealed a disproportionate contribution of hepatic stellate cells to the cellular communication network. Although they comprised only 8% of the normal cell population, hepatic stellate cells were responsible for 48% of the ligands predicted to engage with receptors on melanoma cells (Fig. 2a). Predominantly, these interactions involved signaling through extracellular matrix components such as collagens, laminins, and fibronectin (Fig. 2b,c). These proteins are known to bind with either CD44, a cell-surface glycoprotein, or with integrin heterodimers, both crucial for cell-cell interactions, adhesion, and migration^22,23^. Additionally, a significant interaction was noted between Hepatocyte Growth Factor (HGF) and MET, a tyrosine kinase receptor (Fig. 2d, Supplementary Table 1 with the full list).

**Figure 2.**
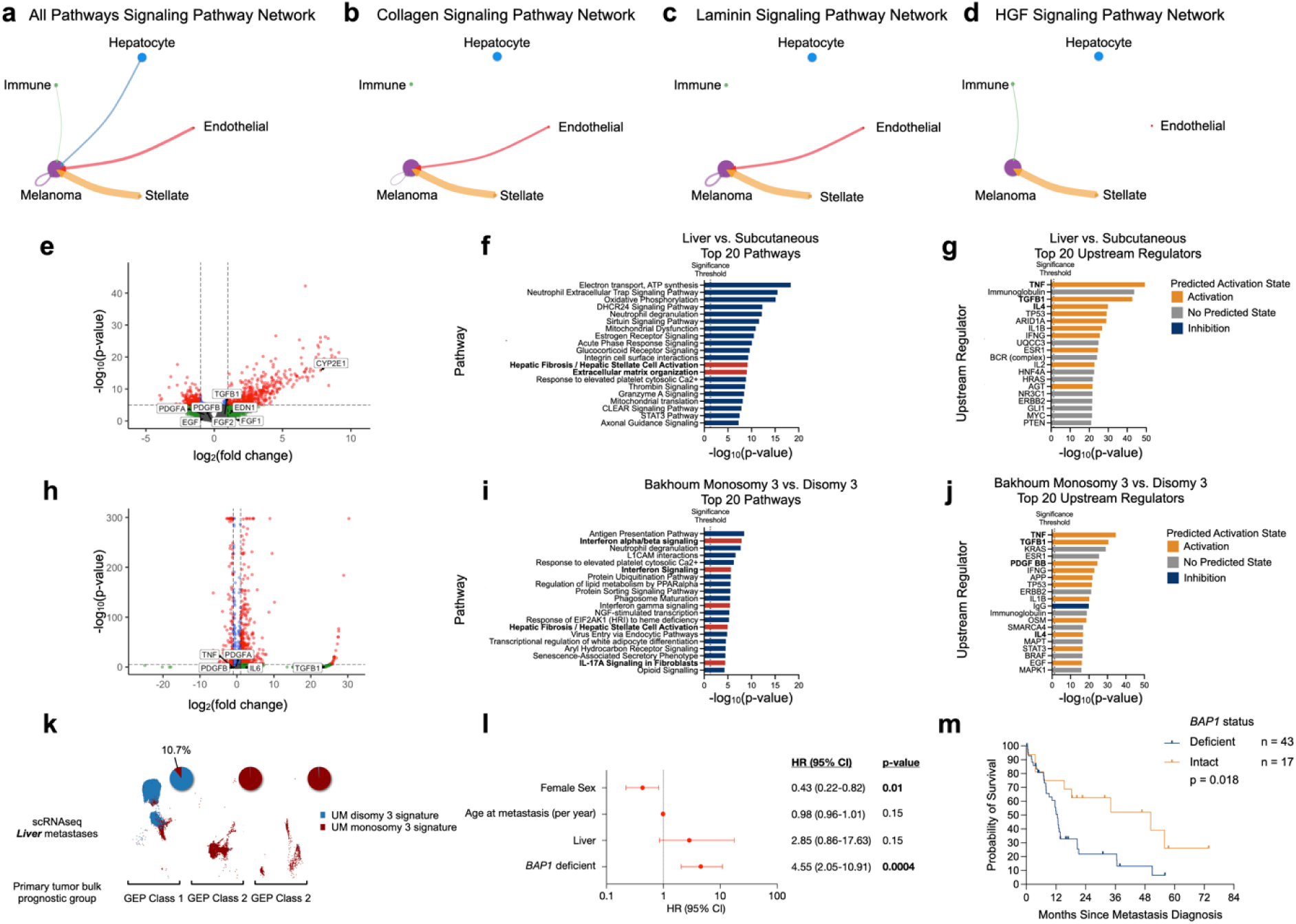
A comprehensive visualization of cell-cell signaling interactions in liver metastases across (**a**) all significant pathways, (**b**) Collagen signaling pathway, (**c**) Laminin signaling pathway, and (**d**) Hepatocyte Growth Factor-MET pathway, where melanoma cells are designated as the target. Interactions are displayed in a circular layout with the area of circles indicating the proportion of cells and thickness of the connecting arrows indicating the strength of the interactions. Pathway analysis of (**e**,**f**,**g**) hepatic vs. extrahepatic metastases and (**h**,**i**,**j**) monosomy 3 vs. disomy 3 UM cells from primary tumors. **e**,**h**, Annotated violin plot of differentially expressed genes between hepatic and extrahepatic metastases, colored by – log_10_(p-value) and log_2_(fold change). Genes related to hepatic stellate cell activation are labeled. Bar plots of top 20 differentially regulated pathways (**f,i**) and upstream regulators (**g**,**j**) using IPA. **k**, UMAP of tumor cells in three hepatic metastases stemming from a GEP1 primary tumor and two GEP2 primary tumors, colored by chromosome 3 signature (disomy shown in blue and monosomy shown in red). **l**, Forest plot for the multivariate Cox proportional hazards regression analysis outcomes, quantifying the impact of several prognostic variables on survival probability in patients with metastatic UM. Hazard ratios (HRs) with 95% confidence intervals (CIs) are shown. The vertical line represents a hazard ratio of 1. **m**, Kaplan-Meier curve showing probability of developing metastasis in patients with primary UM stratified by *BAP1* status (deficient shown in blue and intact shown in orange). Statistical significance was determined using the Log-rank test. *Source: **a**,**b**,**c**,**d**, Durante et al.*^18^*; **e**,**f**,**g**, Karlsson et al.*^12^*; **h**,**i**,**j**, Bakhoum et al.*^7^; ***k****, Wang et al.*^19^*; **l**,**m**, Nguyen et al.*^14^

Recognizing the established role of many of these signaling interactions in driving metastasis across various cancers, we sought to determine whether their upregulation was particularly significant in hepatic versus extrahepatic UM metastases. We compared transcriptional profiles between hepatic (n = 30) and extrahepatic (n = 8, subcutaneous) UM samples^12^. Ingenuity Pathway Analysis (IPA)^24^ and Gene Set Enrichment Analysis (GSEA)^25^ revealed upregulation of pathways and regulators integral to hepatic stellate cell signaling (Fig. 2e,f,g, Extended Data Fig. 3a, and Supplementary Table 2), including hepatic stellate cell activation, extracellular matrix organization, TGF-β, IL6, and PDGFB^26^. These findings emphasize the significance of tumor cell-hepatic stellate cell crosstalk in the development of hepatic UM metastasis.

While hepatic stellate cells constitute only 5-8% of hepatic cells, they have a pivotal role in liver fibrogenesis^27,28^. Activated hepatic stellate cells can foster a pro-inflammatory and pro-angiogenic milieu conducive to metastatic spread in different cancers including UM^29,30^. With the observed inclination of UM clones — specifically those monosomic for chromosome 3 and exhibiting a GEP2 phenotype — to colonize the liver, we examined if these clones express transcriptional programs that are primed to activate stellate cells while still within the primary tumor. For this purpose, we utilized single cell-RNA sequencing data from enucleation specimens of primary UMs (n = 6)^7^ (Extended Data Fig. 4a,b,c), and from a published cohort (n = 8)^18^ (Extended Data Fig. 4d,e,f). Tumor cells were assigned to a prognostic class based on average expression of specific genes derived from TCGA (monosomy 3 vs. disomy 3 tumors) and fitted to a two-component Bayesian Gaussian mixture model (refer to Extended Data Fig. 5a,b and Methods). Differential gene expression analysis between monosomy 3 and disomy 3 primary UM cells demonstrated that monosomy 3 cells displayed expression of genes and pathways known to activate hepatic stellate cells, including TGFB1, VEGF, IL6, and PDGFB (Fig. 2h,i,j, Extended Data Fig. 6a,b,c,d,e, and Supplementary Tables 3 and 4). *In vitro*, primary UM cell lines characterized by monosomy 3 and a GEP class 2 phenotype (Extended Data Fig. 7a)., exhibited upregulation of hepatic stellate cell-activating transcriptional programs (Extended Data Fig. 7b,c,d,e and Supplementary Table 5), in contrast to those with disomy 3 and a GEP class 1 phenotype. These findings suggested that tumor clones with monosomy 3 signature have increased predilection to the liver likely due to their enhanced capacity to activate hepatic stellate cells.

We then interrogated clonal composition in hepatic metastases through the lens of primary UM clonal heterogeneity. We analyzed the profiles of single-cell RNA sequencing of three hepatic UM metastases (Extended Data Figure 2a,b,c), where the GEP of the corresponding primary tumors had been determined through bulk tumor profiling (Extended Data Fig. 5b)^18^. Notably, one of the hepatic metastatic samples originated from an individual whose primary tumor exhibited a GEP class 1 phenotype, in contrast to the other two samples which originated from GEP class 2 primary tumors. This exceptional case — a GEP class 1 primary tumor leading to liver metastasis — prompted us to dissect the clonal composition of the three hepatic metastases. Remarkably, nearly all cells in the two hepatic metastases originating from a GEP class 2 primary exhibited a monosomy 3 signature (Fig. 2k). In contrast, the hepatic specimen originating from a GEP class 1 tumor had mixed clones: 89% disomy 3 and 11% monosomy 3 cells. This finding, albeit in a single case, suggest that the presence of monosomy 3 cells may be necessary for liver metastasis formation, possibly due to their ability to activate hepatic stellate cells.

The predisposition of specific UM clones for liver metastasis prompted us to explore factors that impact survival outcomes in patients with hepatic metastases. We had previously observed that patients with hepatic metastases typically experienced worse survival outcomes than those with solely extrahepatic metastases^6^. However, we now considered whether tumor-specific characteristics could also play a significant role, not just the site of metastasis. To assess the relative impacts of *BAP1* loss and liver involvement on overall survival, we performed a Cox proportional hazard regression analysis using clinical and genomic data obtained from 69 individuals with metastatic UMs that were profiled using whole exome sequencing^14^ (Extended Data Table 3). Within a multivariable model, which encompasses sex, age at metastasis, metastatic site (liver), and *BAP1* status in metastatic specimen as variables, the presence of *BAP1*-deficient clones was linked to reduced overall survival (*P* = 0.0004, HR = 4.55, 95% CI = 2.05-10.91) (Fig. 2l). Interestingly, neither age nor the site of metastasis (liver) significantly influenced overall survival in this model. Female sex correlated with improved overall survival, consistent with prior studies^6^. Further analysis using Kaplan-Meier survival curves specifically for patients with liver-only metastasis, the presence of *BAP1* mutations in metastasis conferred worse survival outcomes (*P* = 0.018) (Fig. 2m). Collectively, these findings suggested that, in addition to their proclivity to spread to the liver, the presence of these clones contributes significantly to the adverse survival outcomes of patients with hepatic metastasis.

## Discussion

Our investigation into the molecular basis of metastatic organotropism in UM provides critical insights into the selective patterns of metastatic dissemination, revealing that this selectivity is not random but driven by specific tumor clones that exist in the primary tumor. This sheds new light on the clonal dynamics of uveal melanoma metastasis, particularly the preferential colonization of the liver. Our analyses reveal that UM clones with certain genetic alterations, notably *BAP1* deficiency and monosomy 3, predispose to liver metastasis and are also associated with poorer patient survival outcomes. This discovery carries significant clinical importance due to the profound impact of hepatic metastases on patient survival.

Our findings highlight the potential for early intervention strategies. We demonstrated that the molecular signature of the primary tumor pre-defines the disease progression pathway. Transcriptional profiling of the primary tumor currently used in clinical practice can, therefore, predict subsequent metastatic patterns. Our results also indicate that adverse survival outcomes are attributed to the intrinsic characteristics of specific tumor clones, beyond the involvement of the liver as a metastatic site. We had recently shown that patients with only extrahepatic metastases had a more favorable prognosis compared to those with hepatic metastases^6^. Here, we show that these adverse survival outcomes stem from the intrinsic characteristics of specific tumor clones, independent of the liver’s involvement as a site of metastasis. Our observations highlight the need to include comprehensive analysis of the metastatic site and genetic variability in ongoing clinical trials.

Furthermore, these results underscore a complex interplay within the tumor-liver microenvironment, presenting potential therapeutic opportunities. We propose a mechanism where tumor clones, particularly those with *BAP1* mutations and monosomy 3, gain an advantage in liver colonization by upregulating transcriptional programs that activate hepatic stellate cells; these programs play a pivotal role in creating a liver environment conducive for tumor cell growth. In the healthy liver, hepatic stellate cells which reside in the liver’s sub-endothelial space of Disse, function to store vitamin A and retinoids^27,28^. However, during liver injury, hepatic stellate cells are activated and produce extracellular matrix^31^. This activation, which is primarily mediated by ligands, including TGFβ, PDGFB, cytokines, and VEGF, enhance cell movement, proliferation, and subsequent production of growth factors and extracellular matrix components^26–28^. Chronically-activated hepatic stellate cells enhance tumor proliferation by forming a scaffold favorable for tumor cells^32^. Coupled with the suppression of immune cells by hepatic stellate cells, this fibrotic setting is highly favorable for tumor growth.

The involvement of hepatic stellate cells in promoting UM metastasis is further validated by immunohistochemistry of metastatic UM samples and *in vitro* studies. Hepatic stellate cells are frequently observed within and surrounding UM metastases^33–36^. Co-culture experiments have highlighted essential interactions between hepatic stellate cells and UM, underscoring the significance of growth factors like FGF9, HGF, and IGF-1, and establishing the role of activated hepatic stellate cells in amplifying the migratory and invasion potential of UM cells^36–38^.

Conversely, UM can stimulate hepatic stellate cells as early as the micro-metastatic phase, and the extracellular vesicles released by UM have been associated with increased hepatic stellate cell responsiveness and contractility^39^. Altogether, our data and these prior studies indicate that the crosstalk between hepatic stellate cells and UM cells are pivotal in establishing the liver-specific tropism in metastatic UM. Our findings provide a hypothesis regarding the role of hepatic stellate cells in promoting hepatic metastasis in UM and highlight the potential utility of HSC-targeted interventions. Therapies that modulate the tumor microenvironment or specifically inhibit hepatic stellate cell activation may present new avenues for treatment.

In conclusion, our findings revealed the molecular mechanisms dictating patterns of metastatic dissemination in UM. This new understanding can improve patient stratification, offers a pathway to tailor therapeutic approaches, and lays the groundwork for the development of novel treatment strategies specifically targeting UM metastasis.

## Methods

### DNA Sequencing and Copy Number Analysis

DNA sequencing data was obtained from 110 hepatic and 34 extrahepatic metastatic specimens to identify hallmark mutations of UM^11–14^. All clinical specimens in this study (n = 5) were collected with informed consent for research use and were approved by the Yale University Institutional Review Boards in accordance with the Declaration of Helsinki. No compensation was provided to participants in this study. Melanoma tumor specimens were excised to alleviate tumor burden, and all analyzed specimens were 2 cm x 2 cm x 2 cm on average and derived from excess surgical material not required for clinical diagnosis and patient care. For data from Nguyen *et al.*, we defined log_2_(fold change) < –0.05 as copy number loss and > 0.05 as copy number gain. *BAP1* intact specimens were defined to have wildtype *BAP1* and disomy 3, while *BAP1* deficient specimens were defined to be *BAP1* mutant and/or have monosomy 3.

### Clinical data and outcomes

All statistical analyses and graphs were generated using GraphPad Prism v9.4.1 (San Diego, California USA). Onset of metastasis Kaplan-Meier curves were generated from 135 patients with primary uveal melanoma treated at Yale New Haven Health (between March 2007 and October 2022) (Fig. 1g,h). We reviewed patients’ medical charts, collecting demographic information including age, sex, race, and ethnicity, in addition to the dates and location of all metastases. The primary outcome was the probability of hepatic or extrahepatic metastasis from time of primary tumor diagnosis. Survivorship Kaplan-Meier curves were generated from Nguyen *et al.* clinical data (n = 69) (Fig. 2m)^14^. The Log-rank method was used to test for significance between groups. Cox proportional hazards regression was used to control for factors (sex, age at metastasis, liver site, and *BAP1* deficiency), and hazards ratios were visualized with a Forest plot.

### scRNA Analysis

Single-cell RNA data of 10 hepatic^18,19^ and 14 primary specimens^7,18^ was analyzed using relevant functions in scanpy^49^. Cells with > 20% of transcripts from mitochondrial genes and low gene counts, defined as greater or less than the median absolute deviation of the total gene counts across all cells, were excluded. Doublets were detected and excluded using sclDblFinder^50^. The filtered raw count matrix was normalized using the shifted logarithm technique. Feature selection was performed to select the top 4,000 deviant genes using the scry package in R^51^. Dimensionality reduction analysis was performed. We clustered cells using the Leiden algorithm and annotated cell types using canonical markers, validating clusters identities using Enrichr (https://maayanlab.cloud/Enrichr/) and WebCSEA (https://bioinfo.uth.edu/webcsea/index.php?csrt=9198285109136139808). Tumor cells were classified based on their expression profiles of genes associated with monosomy and disomy 3 UMs (Supplementary Table 6 provided with genes). As described by Bakhoum *et al.*^7^, we employed a two-component Bayesian Gaussian Mixture Model (GMM), where we fit the model to a two-dimensional data distribution and computed using the diagonal covariance matrix of each component, using the Python package scikit-learn^52^. The model was used to predict the most probably classification of tumor cells into either monosomy 3 or disomy 3 clones. We performed differential gene expression analysis between monosomy 3 vs. disomy 3 cells using the rank_genes_groups function in scanpy based on the Wilcoxon rank sum test. We plotted volcano plots using the EnhancedVolcano package in R^53^.

### CellChat analysis

To predict ligand-receptors interactions, we used the CellChat R package^21^. Single-cell RNA sequencing data were processed using scanpy^49^, and log-normalized counts were exported into CSV format. A CellChat object was created that integrated expression data with corresponding metadata. The identification of cell types was based on manual cell annotations, as detailed in Extended Data Fig. 5a,b. We used the CellChatDB.human database which is tailored for human cell-cell communication studies and focuses on interactions mediated by secreted signaling molecules. We used the netAnalysis function to compute potential cell-cell communication networks based on known ligand-receptor pairs. We utilized various functions within CellChat, such as netVisual_circle and netVisual_bubble, to visualize the number and strength of these interactions.

### Bulk RNA Sequencing Analysis

Raw sequencing reads were aligned to the human reference genome, GRCh38, using the STAR alignment tool in Partek^®^ Flow^®^ (Partek Inc.). For differential gene expression between hepatic and extrahepatic specimens and UM cell lines MP38 vs. 92.1, we utilized the DESeq2 method. For visualization of differential gene expression results using volcano plots, we used the EnhancedVolcano package in R^53^. Genes of interest were highlighted based on their fold change in expression.

### Differential Gene Expression Analysis

For differential gene expression from single-cell RNA sequencing data, we used the FindMarkers function in scanpy^49^ to identify genes differentially expressed between monosomy 3 and disomy 3 UM cells. For visualization of differential gene expression results using volcano plots, we used the EnhancedVolcano package in R^53^. Genes of interest were highlighted based on their statistical significance and their fold change in expression.

### Gene set enrichment analysis

GSEA was performed using the GSEA software (The Broad Institute)^25^ on TMM-normalized bulk RNA-Seq data using the c2.all.v2023.2.Hs.symbols.gmt gene set. We performed GSEA on analyzed Log-normalized scRNA data by running a t-test using rank_genes_groups in scanpy and using the run_gsea function in decoupler on the differential expression scores on the c2.all.v2023.2.Hs.symbols.gmt gene set. Gene signatures were plotted as a function of their NES and –log_10_(FDRq) values (bulk RNAseq) and –log10(p-value) (scRNA). Complete GSEA results are listed in Supplementary Tables 2-5.

### Ingenuity Pathway Analysis

To identify upstream regulators and significant pathways from differential gene expression, we employed Ingenuity Pathway Analysis (IPA, QIAGEN)^24^. IPA is a powerful bioinformatics tool that enables the integration and interpretation of data from gene expression. Significance was determined by a false discovery rate (FDR) threshold of greater than 0.1, a log_2_(fold change) cutoff between –1.5 and 1.5, ensuring that the analysis is focused on genes with the most biological significance. We performed unrestricted analysis, not defining species, cell type, or other characteristics.

The Core Analysis in IPA quickly identifies pathways, relationships, and mechanisms relevant to changes observed in the dataset. The Canonical Pathways analysis identifies canonical pathways across the dataset by considering the activity of one or more key molecules in pathway and their causal relationships with each other. The overall activity states are predicted based on a z-score algorithm. Significance is calculated as the p-value of overlap, using the right-tailed Fisher’s Exact Test.

The Upstream Regulator analysis identifies surface molecules and uses a z-score algorithm the cascade of upstream transcriptional regulators that could be responsible for the observed gene expression changes in the dataset. Regulators with an activation z-score greater or equal to 2 are reported as Activated (orange), while those with an activation z-score less than or equal to – 2 are reported as Inhibited (blue). Regulators for which IPA does not make a prediction are reported as having No Predicted State (gray). P-value of overlap is calculated with a right-tailed Fisher’s Exact test, which unlike the z-score, does not consider gene expression or phosphorylation state.

## Data Availability

Clinical data is available upon request. All patients’ data analyzed from published studies are publicly available accordingly. All raw sequencing data generated for this study will be deposited to NCBI’s Gene Expression Omnibus (GEO) database prior to publication. The Bakhoum *et al.* 2021 bulk RNAseq data is available at GSE181600, and the single cell RNA sequencing data is available under accession code GSE160883. The Durante *et al.* 2020 single cell RNA sequencing data is available in GEO under accession code GSE139829. The Ny *et al.* 2021 data is available in the European Genome-phenome Archive (EGA) under the accession code EGAS00001005478. The Karlsson *et al.* 2020 data used in this study is available in EGA under the accession code EGAS00001004296. The Nguyen *et al.*^14^ data was download from the cBioPortal study MSK-MetTropism (MSK, Cell 2021) and was filtered for patients with metastatic uveal melanoma. The TCGA Uveal melanoma data by Robertson *et al.*^10^ was accessed through cBioPortal (Uveal Melanoma (TCGA, PanCancer Atlas))^40–48^. The Wang *et al.* 2023 data is available in GEO under accession code GSE192402.

## Extended Data Tables

**Extended Data Table 1.** Patient Characteristics, Whole Exome Data. Source: Yale, Karlsson et al. ^12^, Ny et al.^13^, Nguyen et al.^14^, and Royer-Bertrand et al.^11^

| **Characteristic** |  |
| --- | --- |
| Age, median (range) | 68.0 (15.9 – 89.0) |
| Sex, n (%) |  |
| Male | 54 (52.4) |
| Female | 49 (47.6) |
| Race, n (%) |  |
| White | 65 (90.3) |
| Asian | 2 (2.8) |
| Other | 1 (1.4) |
| Not disclosed | 4 (5.5) |
| Lifetime metastasis, n (%) |  |
| Liver only | 30 (29.1) |
| Extrahepatic only | 12 (11.7) |
| Liver and extrahepatic | 61 (59.2) |

**Extended Data Table 2.** Patient Characteristics, Kaplan Meier Survival Analysis. *Source: Yale*

| **Characteristic** |  |
| --- | --- |
| Age, median (range) | 71 (25.0 – 96.0) |
| Gender, n (%) |  |
| Male | 75 (55.6) |
| Female | 60 (44.4) |
| Race, n (%) |  |
| White | 127 (94.1) |
| Asian | 1 (0.7) |
| Other | 6 (4.5) |
| Refused | 1 (0.7) |
| First metastatic episode, n |  |
| Liver | 36 |
| Extrahepatic | 24 |
| GEP class, n (%) |  |
| Class 1 | 89 (65.9) |
| Class 2 | 46 (34.1) |

**Extended Data Table 3.** Patient Characteristics, Multivariate Cox proportional hazards regression. *Source: Nguyen et al.*^14^

| Characteristic |  |
| --- | --- |
| Age, median (range) | (62.4, 15.9-89.0) |
| Sex, n (%) |  |
| Male | 36 (52.2) |
| Female | 33 (47.8) |
| Race, n (%) |  |
| White | 60 (89.5) |
| Asian | 2 (3.0) |
| Other | 1 (1.5) |
| Refused | 4 (6.0) |
| Metastasis, n (%) |  |
| Liver only | 20 (29.0) |
| Extrahepatic only | 9 (13.0) |
| Liver and extrahepatic | 40 (58.0) |

**Extended Data Figure 1.**
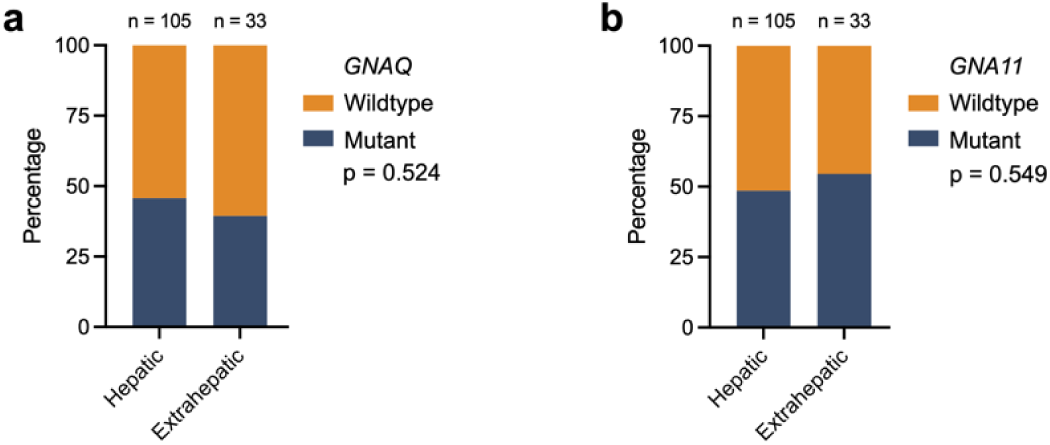
Proportion of *GNAQ* (**a**) and (**b**) *GNA11* mutations (mutant shown in blue and wildtype shown in orange) in hepatic and extrahepatic metastases. Statistical significance was determined using a two-sided Chi-square test. *Source: **a**,**b**, Yale, Ny et al.*^13^*, Karlsson et al.*^12^*, Nguyen et al.*^14^*, and Royer-Bertrand et al.*^11^

**Extended Data Figure 2.**
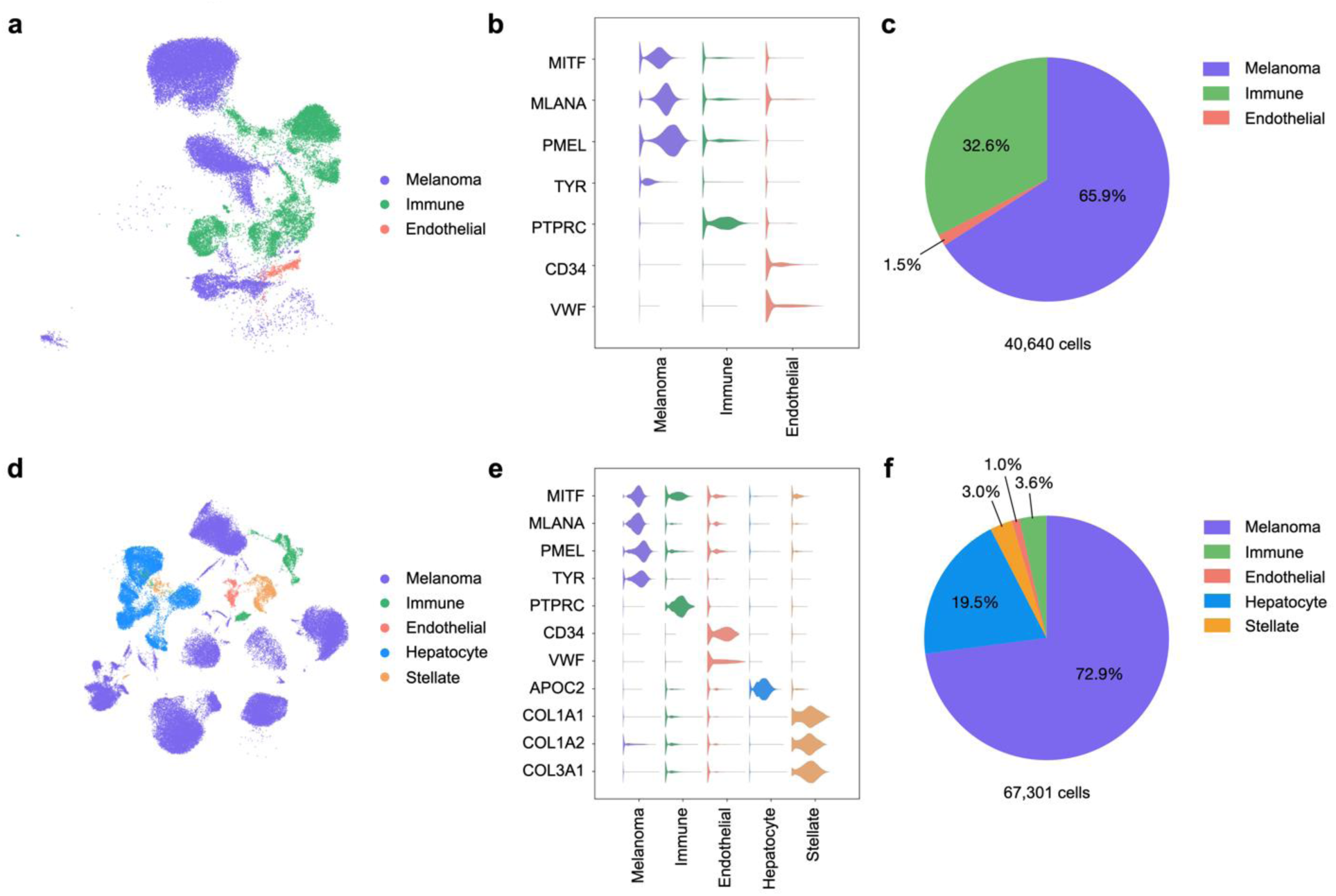
Single-cell data of 3 (**a**,**b**,**c**) and 7 (**d**,**e**,**f**) from hepatic UM metastases. (**a**,**d**) UMAP projections, with individual cells colored by predicted cell types (Melanoma, purple; Immune, green; Endothelial, salmon; Hepatocyte, light blue; Stellate, yellow) (**b**,**e**). Violin plots showing the normalized expression of canonical markers across the different cell types. (**e**,**f**) Pie charts showing the proportion of cell types in each study. *Source: **a**,**b**,**c**, Durante et al.*^18^*; **d**,**e**,**f**, Wang et al.*^19^

**Extended Data Figure 3.**
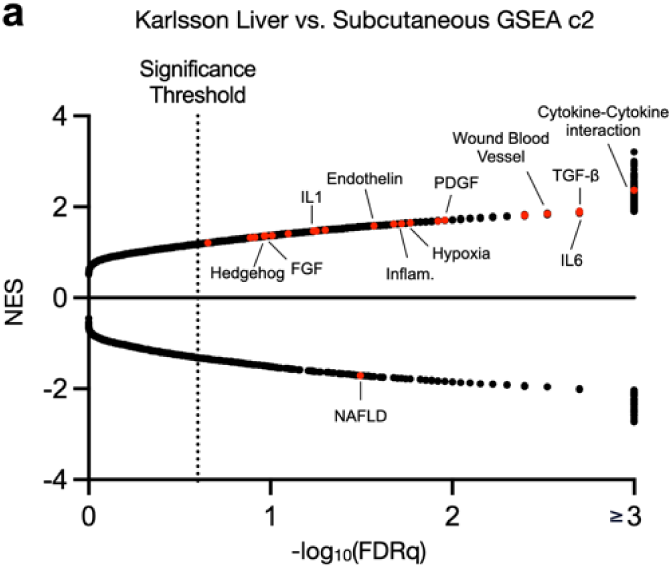
**a**, Volcano plots showing top differentially-regulated genes in hepatic and subcutaneous metastases. Normalized enrichment scores (NES) of gene sets are shown as a function of significance (-log_10_(FDR_q_) or –log_10_(p-value)). Significant pathways related to hepatic stellate cell activation are colored red, while non-significant related pathways are colored in gray. *Source: **a**, Karlsson et al.*^12^

**Extended Data Figure 4.**
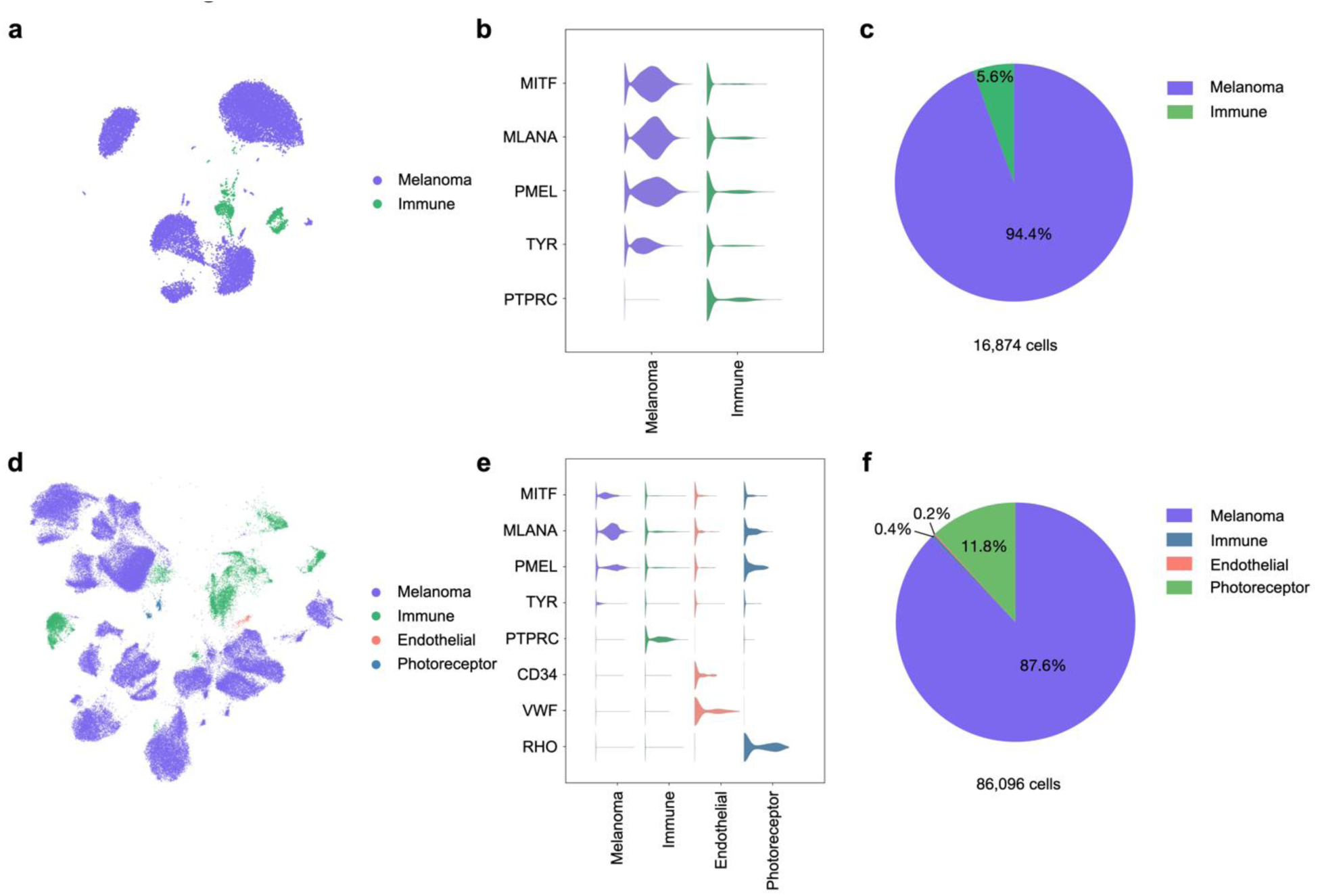
Single-cell data of (**a**,**b**,**c**) 6 and (**d**,**e**,**f**) 8 primary UM enucleation specimens. (**a**,**d**) UMAP projections, with individual cells colored by predicted cell types according to their expression of canonical genes (Melanoma, purple; Immune, green; Endothelial, salmon; Photoreceptor, dark blue). (**b**,**e**) Violin plots showing the normalized expression of canonical markers across the different cell types. (**c**,**f**) Pie charts showing the proportion of cell types in each study. *Source: **a**,**b**,**c**, Bakhoum et al.*^7^*; **d**,**e**,**f**, Durante et al.*^18^

**Extended Data Figure 5.**
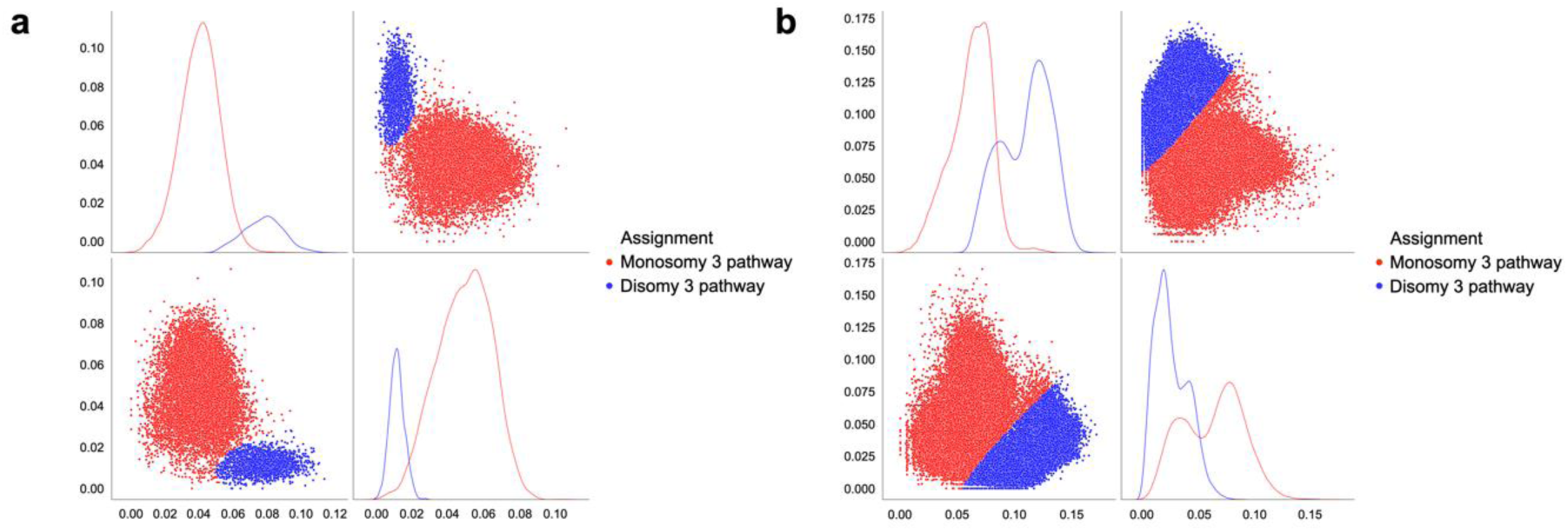
**a**,**b**, Individual tumor cells from primary UMs assigned to monosomy 3 (red) and disomy 3 cells (blue) according to their average expression of monosomy 3 or disomy 3 signatures and by fitting a two-component Bayesian Gaussian Mixture Model as described in the Methods section and in Bakhoum *et al.*^7^. *Source: **a**, Bakhoum et al.*^7^*; **b**, Durante et al.*^18^

**Extended Data Figure 6.**
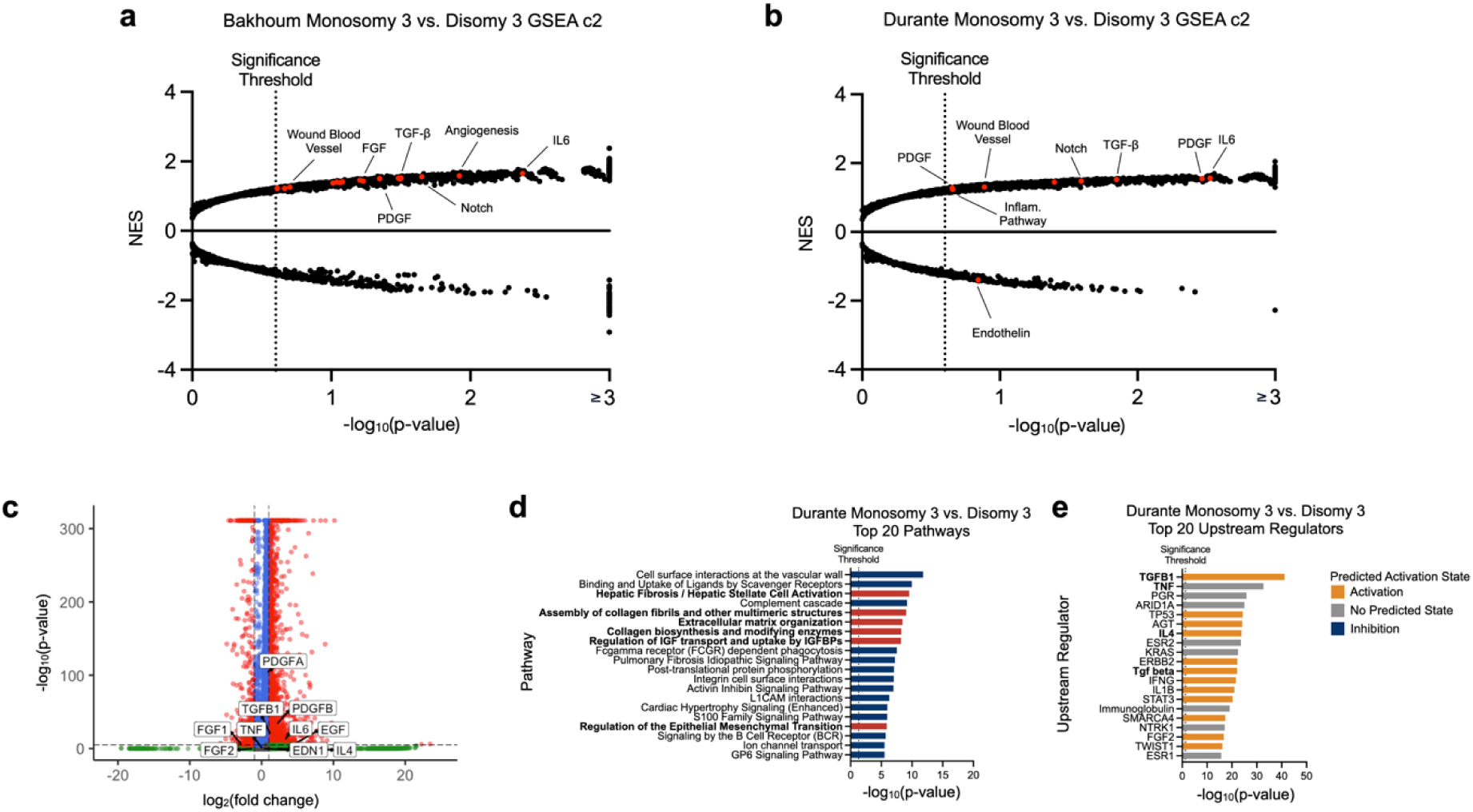
**a**,**b**, Volcano plots showing top differentially-regulated genes in monosomy 3 and disomy 3 primary metastases from two different data sets. Normalized enrichment scores (NES) of gene sets are shown as a function of significance (-log_10_(FDR_q_) or – log_10_(p-value)). Significant pathways related to hepatic stellate cell activation are colored red. **c**, Annotated violin plot of differentially expressed genes between monosomy 3 and disomy 3 and cells in primary UMs analyzed by single-cell RNA sequencing, colored by –log_10_(p-value) and log_2_(fold change). Genes related to hepatic stellate cell activation are labeled. **d**,**e**, Bar plots of top 10 differentially regulated (**d**) pathways and (**e**) upstream regulators between monosomy 3 and disomy 3 clones in primary UMs. Pathways and upstream regulators related to hepatic stellate cell activation are bolded (**d**,**e**) and highlighted in red (**d**). *Source: **a**, Bakhoum et al.*^7^*; **b**,**c**,**d**,**e**, Durante et al.*^18^

**Extended Data Figure 7.**
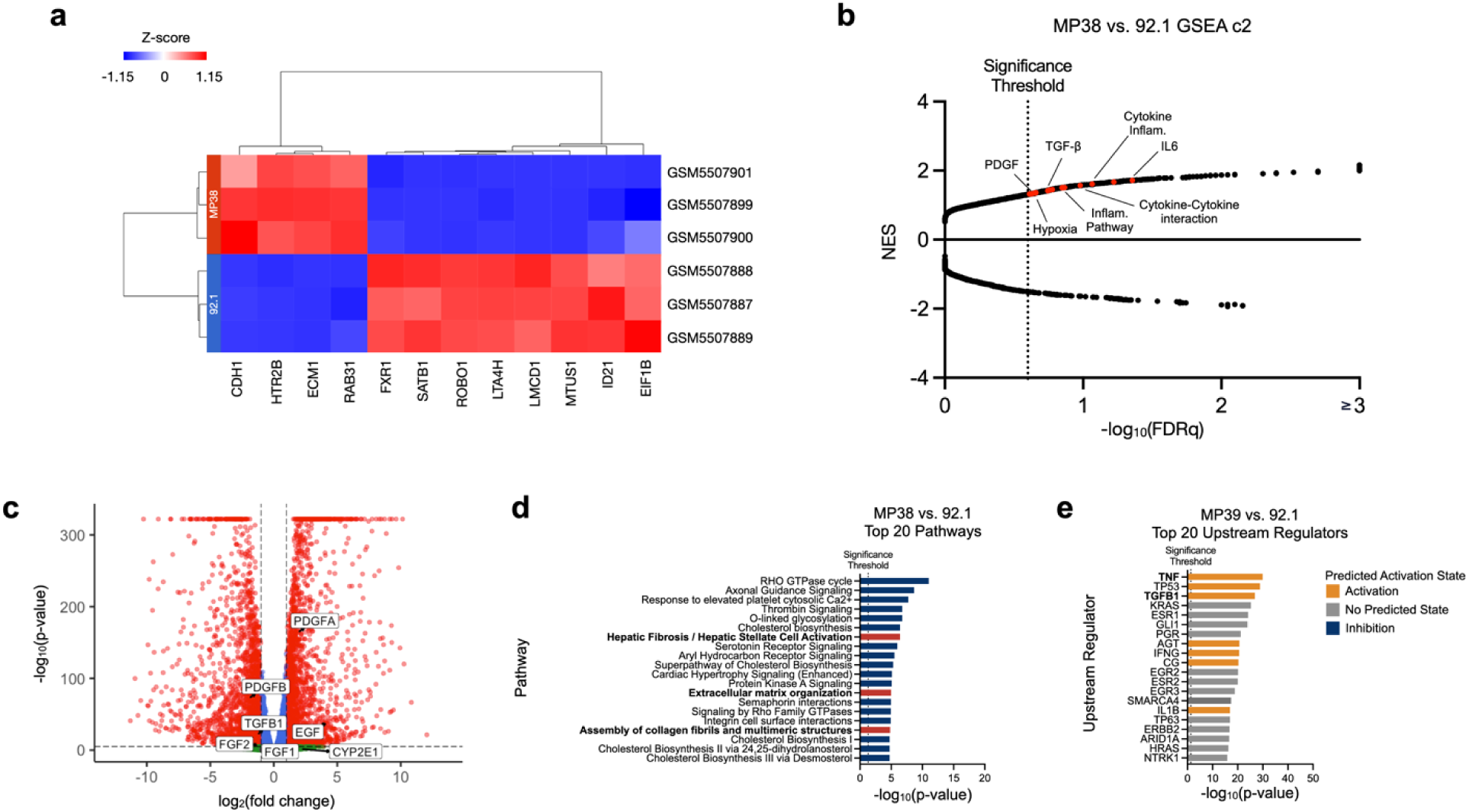
**a**, Unbiased hierarchical clustering of UM cell lines, 92.1 and MP38 across biological triplicates, based on normalized expression values of the 12-discriminant GEP gene set ^7^. **b**, Volcano plots showing top differentially expressed gene sets in UM cells, MP38 vs. 92.1. Normalized enrichment scores (NES) of gene sets are shown as a function of significance (– log_10_(FDR_q_) or –log_10_(p-value)). Significant pathways related to hepatic stellate cell activation are colored red. **c**, Annotated volcano plot of differentially expressed genes between UM cell lines, 92.1 (GEP1) and MP38 (GEP1),^7^ colored by –log_10_(p-value) and log_2_(fold change). Genes related to hepatic stellate cell activation are labeled. **d**,**e**, Bar plots of top 10 differentially regulated (**d**) pathways and upstream regulators between 92.1 and MP38 (bulk RNAseq). Pathways and upstream regulators related to hepatic stellate cell activation are bolded (**d**,**e**) and highlighted in red (**d**). *Source: **a**,**b**,**c**,**d**,**e**, Bakhoum et al.*^7^

## References

1. Jager, M.J., et al. Uveal melanoma. Nat Rev Dis Primers 6, 24 (2020).

2. Gao, Y., et al. Metastasis Organotropism: Redefining the Congenial Soil. Dev Cell 49, 375–391 (2019).

3. Nguyen, D.X., Bos, P.D. & Massague, J. Metastasis: from dissemination to organ-specific colonization. Nat Rev Cancer 9, 274–284 (2009).

4. Collaborative Ocular Melanoma Study, G. Assessment of metastatic disease status at death in 435 patients with large choroidal melanoma in the Collaborative Ocular Melanoma Study (COMS): COMS report no. 15. Arch Ophthalmol 119, 670–676 (2001).

5. Diener-West, M., et al. Development of metastatic disease after enrollment in the COMS trials for treatment of choroidal melanoma: Collaborative Ocular Melanoma Study Group Report No. 26. Arch. Ophthalmol. 123, 1639–1643 (2005).

6. Demkowicz, P., et al. Determinants of overall survival in patients with metastatic uveal melanoma. Cancer 129, 3275–3286 (2023).

7. Bakhoum, M.F., et al. Loss of polycomb repressive complex 1 activity and chromosomal instability drive uveal melanoma progression. Nat Commun 12, 5402 (2021).

8. Bakhoum, M.F. & Esmaeli, B. Molecular Characteristics of Uveal Melanoma: Insights from the Cancer Genome Atlas (TCGA) Project. Cancers (Basel) 11(2019).

9. Robertson, A.G., et al. Comprehensive Molecular Characterization of Muscle-Invasive Bladder Cancer. Cell 171, 540–556 e525 (2017).

10. Robertson, A.G., et al. Integrative Analysis Identifies Four Molecular and Clinical Subsets in Uveal Melanoma. Cancer Cell 32, 204–220 e215 (2017).

11. Royer-Bertrand, B., et al. Comprehensive Genetic Landscape of Uveal Melanoma by Whole-Genome Sequencing. Am J Hum Genet 99, 1190–1198 (2016).

12. Karlsson, J., et al. Molecular profiling of driver events in metastatic uveal melanoma. Nat Commun 11, 1894 (2020).

13. Ny, L., et al. The PEMDAC phase 2 study of pembrolizumab and entinostat in patients with metastatic uveal melanoma. Nat Commun 12, 5155 (2021).

14. Nguyen, B., et al. Genomic characterization of metastatic patterns from prospective clinical sequencing of 25,000 patients. Cell 185, 563–575 e511 (2022).

15. Tschentscher, F., et al. Tumor classification based on gene expression profiling shows that uveal melanomas with and without monosomy 3 represent two distinct entities. Cancer Res. 63, 2578–2584 (2003).

16. Onken, M.D., Worley, L.A., Ehlers, J.P. & Harbour, J.W. Gene expression profiling in uveal melanoma reveals two molecular classes and predicts metastatic death. Cancer Res. 64, 7205–7209 (2004).

17. Prescher, G., et al. Prognostic implications of monosomy 3 in uveal melanoma. Lancet 347, 1222–1225 (1996).

18. Durante, M.A., et al. Single-cell analysis reveals new evolutionary complexity in uveal melanoma. Nat Commun 11, 496 (2020).

19. Wang, Y., et al. Multimodal single-cell and whole-genome sequencing of small, frozen clinical specimens. Nat Genet 55, 19–25 (2023).

20. Andrews, T.S., et al. Single-Cell, Single-Nucleus, and Spatial RNA Sequencing of the Human Liver Identifies Cholangiocyte and Mesenchymal Heterogeneity. Hepatol Commun 6, 821–840 (2022).

21. Jin, S., et al. Inference and analysis of cell-cell communication using CellChat. Nat Commun 12, 1088 (2021).

22. Senbanjo, L.T. & Chellaiah, M.A. CD44: A Multifunctional Cell Surface Adhesion Receptor Is a Regulator of Progression and Metastasis of Cancer Cells. Front Cell Dev Biol 5, 18 (2017).

23. Mezu-Ndubuisi, O.J. & Maheshwari, A. The role of integrins in inflammation and angiogenesis. Pediatr Res 89, 1619–1626 (2021).

24. Kramer, A., Green, J., Pollard, J., Jr. & Tugendreich, S. Causal analysis approaches in Ingenuity Pathway Analysis. Bioinformatics 30, 523–530 (2014).

25. Subramanian, A., et al. Gene set enrichment analysis: a knowledge-based approach for interpreting genome-wide expression profiles. Proc Natl Acad Sci U S A 102, 15545–15550 (2005).

26. Tsuchida, T. & Friedman, S.L. Mechanisms of hepatic stellate cell activation. Nat Rev Gastroenterol Hepatol 14, 397–411 (2017).

27. Yin, C., Evason, K.J., Asahina, K. & Stainier, D.Y. Hepatic stellate cells in liver development, regeneration, and cancer. J Clin Invest 123, 1902–1910 (2013).

28. Hellerbrand, C. Hepatic stellate cells--the pericytes in the liver. Pflugers Arch 465, 775–778 (2013).

29. Babchia, N., Landreville, S., Clement, B., Coulouarn, C. & Mouriaux, F. The bidirectional crosstalk between metastatic uveal melanoma cells and hepatic stellate cells engenders an inflammatory microenvironment. Exp Eye Res 181, 213–222 (2019).

30. Cheng, H., et al. Co-targeting HGF/cMET Signaling with MEK Inhibitors in Metastatic Uveal Melanoma. Mol. Cancer Ther. 16, 516–528 (2017).

31. Vidal-Vanaclocha, F. The prometastatic microenvironment of the liver. Cancer Microenviron 1, 113–129 (2008).

32. Cogliati, B., Yashaswini, C.N., Wang, S., Sia, D. & Friedman, S.L. Friend or foe? The elusive role of hepatic stellate cells in liver cancer. Nat Rev Gastroenterol Hepatol 20, 647–661 (2023).

33. Grossniklaus, H.E. Progression of ocular melanoma metastasis to the liver: the 2012 Zimmerman lecture. JAMA Ophthalmol 131, 462–469 (2013).

34. Krishna, Y., McCarthy, C., Kalirai, H. & Coupland, S.E. Inflammatory cell infiltrates in advanced metastatic uveal melanoma. Hum. Pathol. 66, 159–166 (2017).

35. Piquet, L., et al. Synergic Interactions Between Hepatic Stellate Cells and Uveal Melanoma in Metastatic Growth. Cancers (Basel) 11(2019).

36. Seitz, T., et al. Role of Fibroblast Growth Factors in the Crosstalk of Hepatic Stellate Cells and Uveal Melanoma Cells in the Liver Metastatic Niche. Int J Mol Sci 23(2022).

37. Barisione, G., et al. Potential Role of Soluble c-Met as a New Candidate Biomarker of Metastatic Uveal Melanoma. JAMA Ophthalmol 133, 1013–1021 (2015).

38. Economou, M.A., et al. Receptors for the liver synthesized growth factors IGF-1 and HGF/SF in uveal melanoma: intercorrelation and prognostic implications. Acta Ophthalmol 86 Thesis 4, 20–25 (2008).

39. Piquet, L., et al. Extracellular Vesicles from Ocular Melanoma Have Pro-Fibrotic and Pro-Angiogenic Properties on the Tumor Microenvironment. Cells 11(2022).

40. Hoadley, K.A., et al. Cell-of-Origin Patterns Dominate the Molecular Classification of 10,000 Tumors from 33 Types of Cancer. Cell 173, 291–304 e296 (2018).

41. Ellrott, K., et al. Scalable Open Science Approach for Mutation Calling of Tumor Exomes Using Multiple Genomic Pipelines. Cell Syst 6, 271–281 e277 (2018).

42. Taylor, A.M., et al. Genomic and Functional Approaches to Understanding Cancer Aneuploidy. Cancer Cell 33, 676–689 e673 (2018).

43. Liu, J., et al. An Integrated TCGA Pan-Cancer Clinical Data Resource to Drive High-Quality Survival Outcome Analytics. Cell 173, 400–416 e411 (2018).

44. Sanchez-Vega, F., et al. Oncogenic Signaling Pathways in The Cancer Genome Atlas. Cell 173, 321–337 e310 (2018).

45. Gao, Q., et al. Driver Fusions and Their Implications in the Development and Treatment of Human Cancers. Cell Rep 23, 227–238 e223 (2018).

46. Poore, G.D., et al. Microbiome analyses of blood and tissues suggest cancer diagnostic approach. Nature 579, 567–574 (2020).

47. Ding, L., et al. Perspective on Oncogenic Processes at the End of the Beginning of Cancer Genomics. Cell 173, 305–320 e310 (2018).

48. Bonneville, R., et al. Landscape of Microsatellite Instability Across 39 Cancer Types. JCO Precis Oncol 2017(2017).

49. Wolf, F.A., Angerer, P. & Theis, F.J. SCANPY: large-scale single-cell gene expression data analysis. Genome Biol 19, 15 (2018).

50. Germain, P.L., Lun, A., Garcia Meixide, C., Macnair, W. & Robinson, M.D. Doublet identification in single-cell sequencing data using scDblFinder [version 2; peer review: 2 approved]. F1000Res 10, 979 (2022).

51. Street, K., Townes, F., Risso, D. & Hicks, S. scry: Small-Count Analysis Methods for High-Dimensional Data. (2023).

52. Pedregosa, F., et al. Scikit-learn: Machine Learning in Python. 2825–2830 (2011).

53. Blighe, K., Rana, S. & Lewis, M. EnhancedVolcano: Publication-ready volcano plots with enhanced colouring and labeling. (2023).

